# Vernalisation-induced changes to the Arabidopsis circadian clock require Polycomb Repressive Complex 2 and are FLC-independent

**DOI:** 10.64898/2026.02.24.707649

**Authors:** Stephanie S. I. Williams, Miguel Montez, Emily Edwards, Pirita Paajanen, Antony N. Dodd

## Abstract

In many plants, prolonged winter cold and seasonal day-length changes align the transition to flowering with spring. This occurs through the vernalisation and photoperiod pathways, respectively. Despite roles for the circadian clock in regulating both pathways, their mechanisms have mainly been studied in isolation and their interactions are not fully understood. We found that vernalisation elicits changes in the circadian clock and this is linked to alterations in several clock-controlled outputs, including photoperiodic flowering. Importantly, vernalisation-induced changes in specific clock genes are stable and persist upon return to warmth, providing the circadian clock with a memory of prior long-term cold exposure. The changes are not systematic and affect specific circadian oscillator genes. This change requires the epigenetic regulator Polycomb Repressive Complex 2 (PRC2), but not the major genetic determinants of vernalisation *FRIGIDA* (*FRI*) and *FLOWERING LOCUS C* (*FLC*). In contrast to their role in *FLC* silencing, core PRC2 components but not accessory proteins are required for these changes in the clock post-vernalisation. Our work raises the possibility that long-term cold feeds epigenetic information into the clock as a seasonal response mechanism, potentially preparing plants for the coming season.

## Introduction

Plants monitor seasonal progression in two broad ways: by tracking day-length changes, and through long-term temperature sensing. In flowering plants, these processes align the transition to reproductive development with the seasonal environment. Some Arabidopsis accessions and winter annual crops require prolonged cold to induce flowering (Bastow et al., 2004; Aikawa et al., 2010). This is known as vernalisation and captures long-term information on past environmental conditions. Species including Arabidopsis and rice are also photoperiodic, and initiate flowering in response to seasonal day length changes (Yanovsky & Kay, 2002; Serrano-Bueno et al., 2017). Photoperiodic responses involve the circadian clock, so the interaction between 24 h circadian timing and long-term environmental monitoring forms an important part of plant phenology. As vernalisation is often studied in isolation from photoperiodism and circadian regulation, we wished to investigate how these processes might be linked in controlling long-term developmental processes.

Circadian clocks adapt plants to environmental fluctuations over both 24 h and annual timescales (Eriksson & Millar, 2003; Millar, 2016). They involve self-sustaining cellular oscillators that are entrained to rhythmic environmental cues such as light and temperature. In turn, the oscillator aligns biological processes with the time of day (Pittendrigh, 1960; Millar, 2003; Hsu & Harmer, 2014). In *Arabidopsis thaliana*, the circadian oscillator comprises a series of interlocking transcription-translation feedback loops. These operate alongside post-transcriptional regulation and chromatin modifications (Fujiwara et al., 2008; Jordi Malapeira et al., 2012; Yang et al., 2020), with histone modifications contributing to the rhythms of some oscillator components (Perales & Mas, 2007; J. Malapeira et al., 2012; Song & Noh, 2012; Baerenfaller et al., 2016). The Arabidopsis circadian clock regulates at least 30% of the transcriptome, suggesting that many processes – including development, physiology, and metabolism – are under circadian regulation (Harmer et al., 2000; Covington et al., 2008; Michael et al., 2008).

In addition to photoperiodism, for some plants the floral transition also requires vernalisation. Vernalisation is thought to prevent premature flowering in autumn when photoperiods may induce flowering, but environmental conditions do not favour reproduction (Hepworth et al., 2018). In Arabidopsis, *FLOWERING LOCUS C* (*FLC*) and *FRIDGIDA* (*FRI)* are factors that specify the vernalisation requirement (Koornneef et al., 1994; Michaels & Amasino, 1999; Sheldon et al., 1999). *FLC* encodes a MADS-box DNA binding transcription factor which, prior to vernalisation, is expressed highly in an open chromatin state, allowing its active transcription. High FLC levels inhibit floral integrator genes including *FT* and *SOC1*, preventing flowering (Michaels & Amasino, 1999). *FRI* is a transcriptional activator of *FLC* and its activity is antagonised by vernalisation (Choi et al., 2011). The Col-0 accession of Arabidopsis has a naturally occurring mutation to the *FRI* allele, rendering it non-functional. Therefore, Col-0 does not require vernalisation and is described as a rapid-cycling accession (Gazzani et al., 2003; Michaels et al., 2003).

During the first stages of vernalisation, transient cold temperatures promote transcriptional *FLC* shutdown through activity of the long noncoding RNA *COOLAIR* (Csorba et al., 2014). Longer cold periods upregulate expression of the PHD finger protein VERNALISATION INSENSITIVE 3 (VIN3) (Sung & Amasino, 2004; Hepworth et al., 2018). VIN3 and its homolog VERNALISATION 5 (VRN5) act as accessory proteins to the Polycomb Repressive Complex 2 (PRC2) (Greb et al., 2007), and are both required for the *FLC* silencing during vernalisation. PRC2 is an evolutionarily conserved chromatin-modifying complex with methyltransferase activity (De Lucia et al., 2008; Hennig & Derkacheva, 2009), which mediates a switch from an active to a repressive chromatin state at *FLC* marked by histone H3 trimethylation at lysine 27 (H3K27me3). Upon the arrival of warmer spring temperatures, PRC2 spreads this repressive mark across the *FLC* locus, conferring stable epigenetic silencing and derepressing flowering (Angel et al., 2011; Yang et al., 2017). *FLC* expression levels are reset for the next generation during embryogenesis (Sheldon et al., 2008; Choi et al., 2009).

There are interactions between the circadian clock and vernalisation pathway. The clock regulates *VIN3* expression during mild low temperatures, such that *VIN3* transcript levels peak at the end of short days (Hepworth et al., 2018). This likely occurs through interaction of the circadian oscillator components CIRCADIAN CLOCK ASSOCIATED1 (CCA1) and LATE ELONGATED HYPOCOTYL (LHY) with the vernalisation-responsive *cis*-element within the *VIN3* promoter (Kyung et al., 2022). Vernalisation is compromised in the *cca1 lhy* double mutant due to impaired upregulation of *VIN3* at 12°C (Kyung et al., 2022). Vernalisation also shortens the circadian period independently from *FLC* expression (Salathia et al., 2006). Furthermore, *FLC* contributes to temperature compensation of the circadian clock, perhaps through interactions with LUX ARRHYTHMO (LUX) and another unknown factor (Edwards et al., 2006). *FLC* also binds weakly to the *LHY* promoter, with this being detectable in *FLC*-overexpressing plants (Spensley et al., 2009). Together, this suggests crosstalk between the vernalisation pathway, *FLC*, and the circadian clock, which might provide an additional mechanism to distinguish between similar photoperiods during autumn and spring (Salathia et al., 2006).

Despite this evidence for interactions between the circadian clock and vernalisation pathway, it remains unclear how vernalisation modulates the circadian oscillator and whether such changes impact plant physiology and development. Therefore, we investigated how specific oscillator components respond to vernalisation, the associated mechanisms within the vernalisation pathway, and outcomes for both growth and photoperiodic flowering responses.

## Results

### Vernalisation induces stable alterations to the circadian clock

We wished to determine whether circadian oscillator dynamics are altered by long-term cold exposure that causes vernalisation. Specifically, we tested whether the oscillator might be altered after return to control temperatures, following vernalisation. We investigated this with the Col *FRI^SF2^* Arabidopsis genotype (henceforth Col *FRI*), a winter annual derivative of Col-0 that contains a functional *FRI* allele introgressed from the San feliu-2 accession, and thus requires vernalisation to flower (Lee et al., 1994). We reasoned that its responses to prolonged cold may differ from those of summer annual (rapid-cycling) Arabidopsis accessions such as Col-0. Moreover, Col *FRI* expresses *FLC* highly before vernalisation (Sung & Amasino, 2004), allowing investigation of the relationship between *FLC* expression and the circadian clock.

Seedlings were cultivated under control temperature conditions (19 °C) for 11 days, and subsequently transferred to vernalising conditions (5 °C, 8 h photoperiod) for 8 weeks (Fig. 1A). This is sufficient to fully silence *FLC* expression (Whittaker & Dean, 2017). Next, the plants were returned to control temperature conditions (19 °C) for 7 days, and subsequently transferred to constant light (LL) to monitor the circadian clock under free running conditions (Fig. 1A). This design was to avoid transient effects within the oscillator that might occur immediately after transfer from the vernalisation treatment to control temperature conditions. Under these control temperature free running conditions, we harvested aerial tissue for mRNA at 4 h intervals over 48 h to examine the dynamics of circadian oscillator transcripts. We compared this with control plants that did not receive a vernalisation treatment. To control for the effect of aging on the circadian clock (Kim et al., 2016), these non-vernalised (NV) plants were grown for an equivalent period of developmental time (26 days) under control temperature conditions (Zhao et al., 2020). Given that the Arabidopsis circadian oscillator is formed from multiple interlocking transcription-translation feedback loops we examined the abundance of several transcripts that function within the oscillator (*CCA1*, *PSEUDO-RESPONSE REGULATOR7* (*PRR7*), *TIMING OF CAB EXPRESSION1* (*TOC1*), *EARLY FLOWERING3* (*ELF3*) and *LUX*). We also evaluated whether transcript dynamics differed between Col-0 and Col *FRI* in NV control plants. In these NV plants, when *FLC* is expressed highly in Col *FRI,* we observed no obvious differences in dynamics of oscillator transcripts between genotypes (Supplementary Fig. 1). *FLC* levels were measured after the 8-week cold treatment; *FLC* transcripts were nearly undetectable after return to control temperatures, demonstrating successful vernalisation (Fig. 1B).

**Figure 1:**
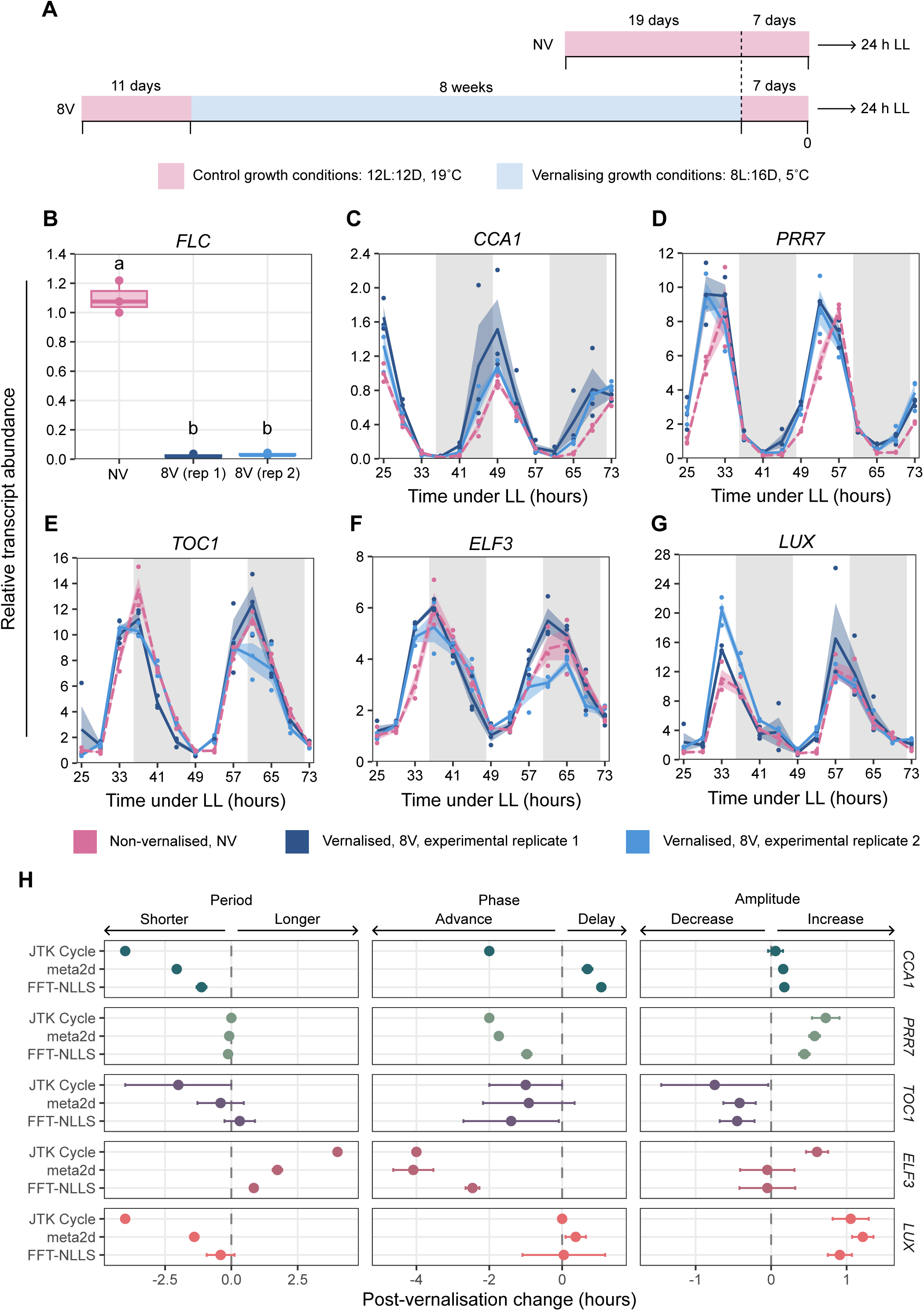
Exposure of Col *FRI* to prolonged cold affects the transcriptional dynamics of circadian oscillator components under subsequent control conditions. **(A)** Arabidopsis Col *FRI* seedlings were initially grown under control temperature conditions (12L:12D 19 °C) for 11 days before transfer to vernalising cold temperatures (8L:16D, 5 °C) for 8 weeks (8V). Following this, plants were returned to 12L:12D 19 °C for a further 7 days. On day 8, plants were transferred to constant light (LL) conditions. Sampling of aerial tissue for RNA started after 25 h of constant light with tissue collected every 4 hours for 48 hours. Non-vernalised (NV) plants were grown for an equivalent developmental time under control temperature conditions (19 °C). For each timepoint, aerial tissue from up to five individual plants was pooled to generate each biological replicate. **(B)** *FLC* transcript levels in Col *FRI* following vernalisation, compared with non-vernalised control plants. Different letters indicate statistically significant differences (p < 0.05), from one-way ANOVA followed by with post hoc Tukey’s HSD test. **(C-G)** Relative abundance of circadian oscillator transcripts *CCA1*, *PRR7*, *TOC1*, *ELF3* and *LUX*. Solid lines indicate mean, with shaded ribbon ± s.e.m (n = 3). Grey shading indicates subjective night. **(H)** The magnitude and change in direction of free running period, phase and amplitude of circadian oscillator transcripts in Col *FRI*, when comparing vernalised and non-vernalised plants. Direction of post-vernalisation change calculated by subtracting the parameter from vernalised samples from the parameter from non-vernalised samples. Point indicates mean difference ± s.e.m (n = 2). Each row represents a different analysis method. In **(B-G)**, NV = non-vernalised, 8V = 8 weeks vernalised. Transcript levels expressed relative to *PP2AA3*, as determined by RT-qPCR.

8 weeks of vernalisation (8V) did not substantially alter the dynamics of dawn-expressed *CCA1* upon return to control temperature conditions. In vernalised and NV plants, maximum *CCA1* transcript levels occurred around subjective dawn (timepoints 25 h and 49 h) and minimum transcript levels around subjective dusk (37 h and 61 h) (Fig. 1C). In contrast, oscillations of transcripts encoding the circadian clock component *PRR7* were altered by the prolonged cold treatment (Fig, 1d). In 8V plants, *PRR7* transcript levels increased earlier during each subjective day compared with the NV plants (29 h and 53 h, *vs*. 33 h and 57 h) (Fig. 1D). The transcripts reached minimum levels at approximately the same time (Fig.1D). Oscillations of transcripts encoding the clock component *TOC1* were not altered consistently between 8V plants and NV plants (Fig. 1E).

*ELF3* and *LUX* encode parts of the evening complex (EC) of the circadian oscillator, and play roles in plant temperature sensing (Nusinow et al., 2011; Silva et al., 2020). In NV plants, the greatest and lowest *ELF3* transcript levels occurred around subjective dusk and subjective dawn, respectively. *ELF3* transcript levels appeared to increase earlier in 8V compared to the NV control, but this was somewhat inconsistent between experimental repeats (Fig. 1F). Our data suggest that *LUX* transcript oscillations were unaltered by the 8-week vernalisation treatment (Fig. 1G).

Using rhythmicity detection algorithms, we identified that all transcripts examined were significantly rhythmic (Supplementary Dataset 1). We detected no systematic change across all oscillator transcripts in their period length, phase, or amplitude following vernalisation (Fig. 1H; Supplementary Dataset 1). However, this analysis identified that both *PRR7* and *ELF3* transcripts phased earlier in 8V plants compared with NV controls (Fig. 1H; Supplementary Dataset 1). While the period of *ELF3* may be lengthened by vernalisation, the period of *PRR7* was unaltered and remained *circa* 24 hours. Moreover, this analysis suggests that exposure to prolonged cold increases the overall amplitude of the rhythm of *PRR7* (Fig. 1H; Supplementary Dataset 1). Using CircaCompare, a tool for statistical comparison of rhythmic datasets (Parsons et al., 2020), we verified that *PRR7* transcripts phase earlier in plants exposed to prolonged cold compared with NV plants (p = 0.015). Overall, this suggests that following a period of prolonged cold, the timing of maximum *PRR7* transcript levels is altered under constant conditions, and this was consistent across both days of experimentation. This could suggest that the repression of *PRR7* is lifted earlier during the subjective day in 8V compared with NV plants.

### Vernalisation alters rhythms of clock output transcripts

Next, we investigated whether vernalisation-induced changes in oscillator dynamics extend to circadian regulated outputs. We quantified the abundance of several transcripts involved in flowering and growth, two major seasonal processes under circadian regulation, and several PRR7-regulated transcripts (Liu et al., 2013). Differences between the accumulation of these transcripts in vernalised and non-vernalised plants were assessed over 24 h of constant light (Fig. 2).

**Figure 2.**
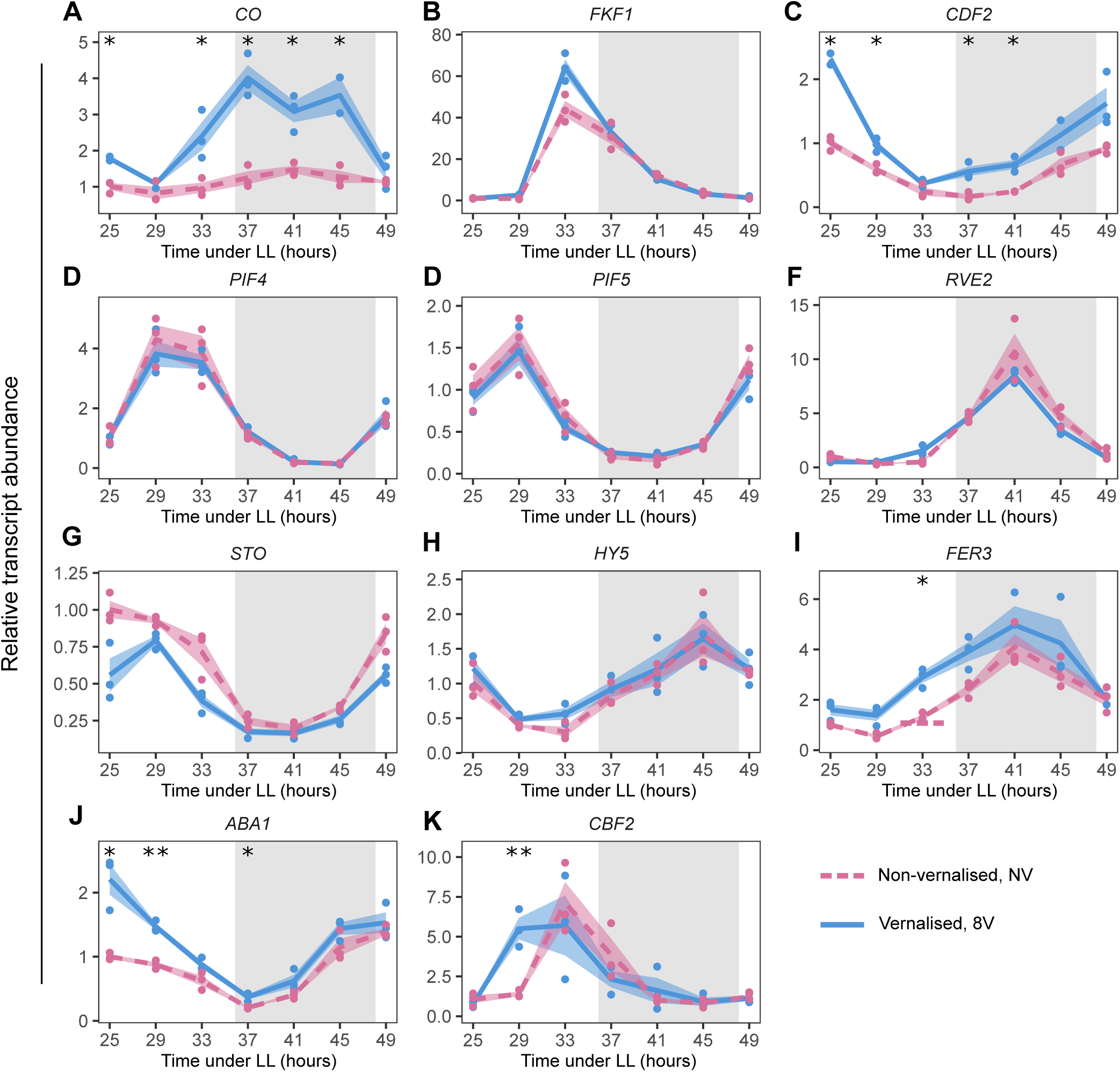
Vernalisation alters the dynamics of transcripts that are outputs from the circadian clock. The abundance of transcripts downstream of the circadian clock that are associated with **(A-C)** flowering time, **(D-F)** growth, and are **(G-K)** known downstream targets of *PRR7* were compared between non-vernalised plants (NV) and those given an 8-week vernalisation treatment (8V) then returned to control conditions and measured under constant light (LL). Transcript levels expressed relative to *PP2AA3* and *UBQ10*, as measured by RT-qPCR. Grey shading represents subjective night. Data are mean ± s.e.m (n = 2-3). Asterisks represent significant differences between vernalised and non-vernalised plants determined by Welch’s two-sample t-tests, p-values adjusted for multiple testing using the Benjamini–Hochberg false discovery rate (FDR) procedure, * p < 0.05, ** p < 0.01.

The circadian clock underpins the regulation of photoperiodic flowering by modulating the timing of peak *CONSTANS* (*CO*) expression. *CO* is a B-box-type zinc-finger transcription factor that directly promotes the floral transition through transcriptional activation of *FLOWERING LOCUS T* (*FT*) (Song et al., 2015). *CO* transcripts were upregulated at the end of the day and during the subjective night in vernalised plants (Fig. 2A). Rhythmic *CO* expression is partly supported by PRR oscillator components (Nakamichi et al., 2007; Hayama et al., 2017) and the CYCLING DOF FACTOR (CDF1-5) protein family (Fornara et al., 2009). Transcripts encoding the blue-light photoreceptor FLAVIN-BINDING KELCH REPEAT F-BOX 1 (FKF1), which mediates degradation of CDF proteins (Sawa et al., 2007), was unaltered following vernalisation (Fig. 2B); however, *CDF2* transcripts were upregulated at some timepoints (Fig. 2C). The circadian clock also exerts control over vegetative growth. PHYTOCHROME INTERACTING FACTORs (PIFs) act downstream of the clock to regulate growth and shade-avoidance responses. *PIF4* and *PIF5* transcripts are regulated by *PRR7* and the EC (Nusinow et al., 2011; Zhang et al., 2020), and transcript levels were unaltered following vernalisation (Fig. 2D, E). Similarly, there were no vernalisation induced changes in *RVE2* transcript levels; RVE2 is also involved in hypocotyl elongation and flowering (Fig. 2F).

*SALT TOLERANCE* (*STO,* known also as *BBX24*) and *ELONGATED HYPOCOTYL 5* (*HY5)* participate in light signalling and photomorphogenesis (Indorf et al., 2007) (Catalá et al., 2011). These are repressed by PRR7 (Liu et al., 2013) but were unaltered following vernalisation (Fig. 2G, H). Expression of iron (Fe) homeostasis genes are also regulated by the circadian clock (Hong et al., 2013). Specifically, the *FERRITINS FER1*, *FER3,* and *FER4* are PRR7-repressed (Liu et al., 2013); we identified an increase in *FER3* transcript levels following vernalisation at 33 h of constant light (Fig. 2I).

The circadian clock influences endogenous ABA levels and the expression of genes involved in its biosynthesis (Covington et al., 2008; Liu et al., 2013). Moreover, over a third of PRR7 targets contain ABA-responsive elements. Following vernalisation, the abundance of *ABA DEFICIENT1* (*ABA1*) transcripts increased around the time of its peak expression (25 h and 29 h, Fig 2. j), suggesting a possible increase in oscillatory amplitude.

*C-REPEAT/DRE BINDING FACTORs* (*CBFs*) are crucial for cold acclimation and are induced rapidly by low temperatures (Fowler et al., 2005). *PRR7* can regulate *CBF*s by binding to their promoters to repress expression around dusk (Liu et al., 2013; Kim et al., 2024). Following vernalisation, *CBF2* transcript abundance was greater in vernalised plants, specifically at 29 hours of constant light (Fig. 2K).

### Vernalisation attenuates photoperiod-regulated differences in flowering time

In Arabidopsis, both vernalisation and photoperiod length regulate the transition to flowering. Arabidopsis is a long-day plant, so flowering is accelerated by longer photoperiods (e.g., 16 h). The circadian clock regulates photoperiodic flowering through stabilisation of CO protein at specific times during the day (Nakamichi et al., 2007; Andrés & Coupland, 2012). Given that the expression of specific circadian clock transcripts was altered following vernalisation (Fig. 1) and *CO* mRNA was upregulated in vernalised plants (Fig. 2A), we hypothesised that this might change the critical daylength threshold for photoperiodic floral induction. To investigate this, following initial growth under short days, we exposed Arabidopsis plants to vernalising long-term cold treatments of two different durations (4 weeks, 8 weeks) alongside a non-vernalised control, and then transferred the plants from all three conditions into either short (8 h light / 16 h dark), equinox (12L:12D), or long (16L:8D) photoperiods under control temperature conditions (19 °C) (Fig. 3A). To understand how the duration of vernalisation treatment and subsequent photoperiod impacted the developmental stage of the transition to flowering, we compared the total leaf number at time of inflorescence emergence.

**Figure 3:**
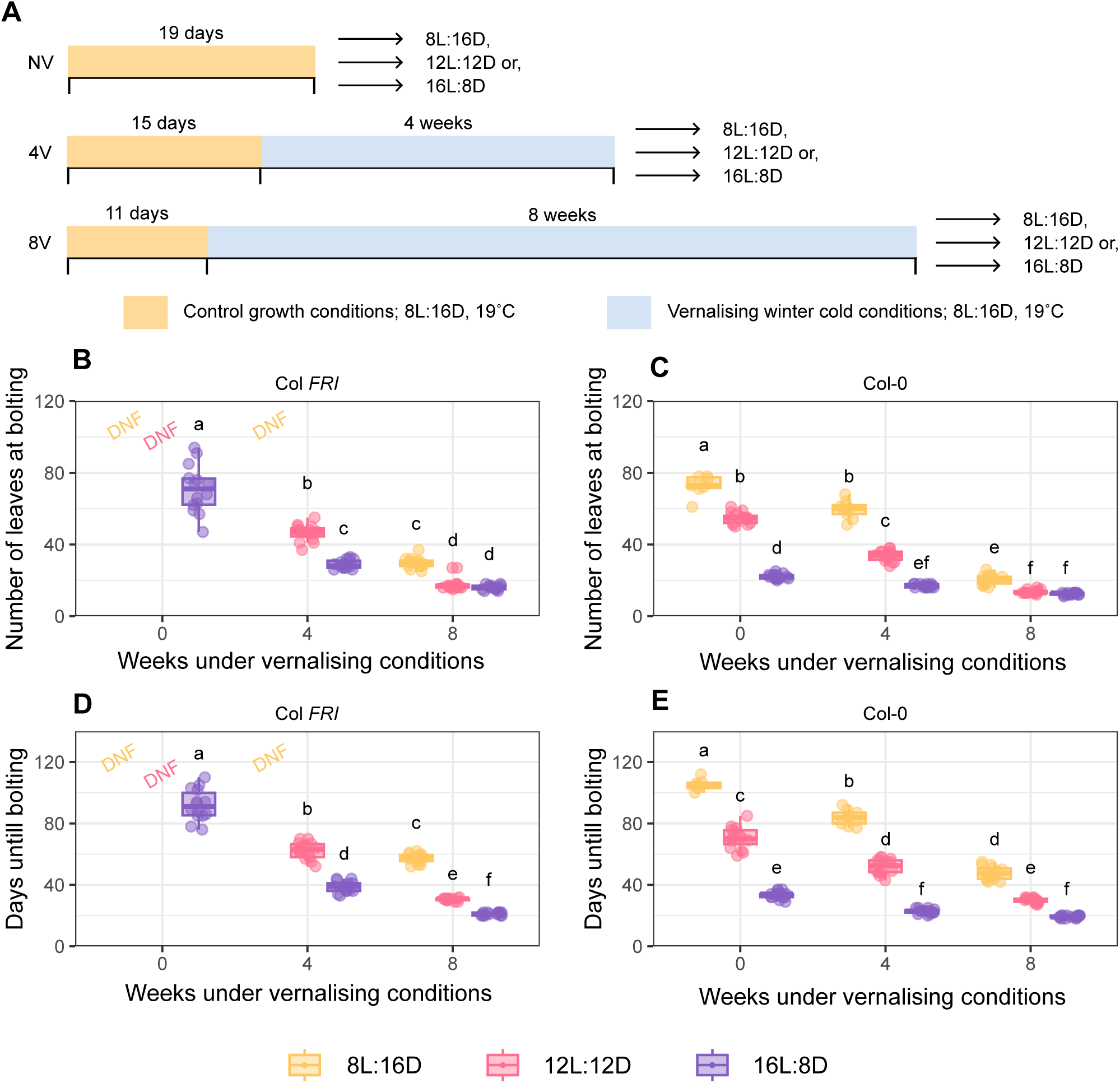
Vernalisation eliminates flowering time differences between equinox and long day conditions**. (A)** Arabidopsis seedlings of Col *FRI* and Col-0 were initially grown under control temperature conditions (8L:16D 19 °C) for at least 11 days before transfer to vernalising cold temperatures (8L:16D, 5 °C) for up to 8 weeks. Differences in cultivation time under control conditions ensured plants were at the same developmental stage after cold treatment. Plants were subsequently grown on compost under control temperatures (19°C) under either 8L:16D, 12L:12D or 16L:8D photoperiods. **(B, C)** Total leaf number at flowering and **(D, E)** days until bolting under short day (8L:16D), equinox (12L:12D), or long day (16L:8D) conditions following either 0, 4, or 8 weeks of vernalising cold in Col *FRI* and Col-0. Groups in which no plant flowered during the timeframe of the experiment are indicated as DNF (did not flower). Different letters indicate statistically significant differences (p < 0.05), from two-way ANOVA followed by with post hoc Tukey’s HSD test, p < 0.05. NV = not vernalised; 4V = 4 weeks vernalisation; 8V = 8 weeks vernalisation.

Longer vernalisation treatments advanced the developmental stage at which flowering occurred (Fig. 3B). This acceleration of flowering is consistent with the well-established vernalisation response of Col *FRI*. Exposure to prolonged cold also caused Col-0 to flower at an earlier developmental stage (Fig. 3C). This was conserved across all photoperiods tested; even under short day conditions, flowering in Col-0 was accelerated by both 4-and 8-week vernalisation treatments relative to non-vernalised plants (Fig. 3C, D). In Col-0 under 16 h photoperiods, there was no further acceleration of flowering when vernalisation was extended from 4 to 8 weeks. This is because plants flowered with the same number of leaves and after the same number of days after both durations of vernalisation (Fig. 3C, E).

Interestingly, exposure to an 8-week vernalisation treatment altered the importance of photoperiod for flowering in both Col-0 and Col *FRI* (Fig. 3B, C). When plants received either no vernalisation treatment or 4 weeks of cold, there was a photoperiod-dependent effect on flowering. Under longer photoperiods, plants flowered at an earlier developmental stage, with 8 h and 12 h photoperiods delaying flowering significantly (Fig. 3B, C). This response was lost after 8 weeks of cold exposure and there was no difference between the total leaf number at bolting between 12 h and 16 h photoperiods in both genotypes (Fig. 3B, C). In Col *FRI*, only 8 weeks of cold was sufficient for flowering under short days within the duration of the experiment (Fig. 3B).

Together, this suggests that exposure of Arabidopsis to prolonged cold for 8 weeks may override the photoperiodic requirement for flowering, allowing plants flower at an earlier developmental stage. Furthermore, this response may be conserved regardless of the genotype’s vernalisation requirement.

### Long-term cold alters growth rates upon return to warmth

The circadian oscillator is associated with vegetative growth regulation of Arabidopsis, and changes in oscillator function can alter growth rates (Dodd et al., 2005; Graf et al., 2010; Chew et al., 2022). Because we observed vernalisation-induced changes in circadian oscillator components and some output transcripts, we hypothesized that there might be changes in growth after long-term cold. To investigate this, we used an experimental design whereby seedlings were vernalised, returned to control temperature conditions, and then transplanted to compost to monitor the visible leaf area of the whole rosette as a proxy for growth. We also compared the growth rate of Col *FRI* with Col-0, to determine whether winter annual accessions might have different growth rates compared with rapidly cycling summer annuals after a period of cold.

To examine differences in plant size we fitted a linear mixed model (LMM) using genotype, treatment, and time as fixed effects, and this also accounted for repeated measures of individual plants. We observed an increase in leaf area with time (p < 0.001), reflecting growth of all plants under control temperatures. This also identified a three-way interaction between treatment, genotype, and time (p < 0.01), suggesting that the effect of winter cold on growth is genotype dependent. Post-hoc testing confirmed that under control temperature conditions, there was no differences in overall rosette area between Col *FRI* or Col-0 plants (p = 0.50). However, whilst vernalisation did not alter the rosette size of Col *FRI* (Fig. 4A, p = 0.86), it did reduce the visible leaf area of Col-0 plants (Fig. 4B, p < 0.01) compared to non-vernalised controls (Fig. 4A-C).

**Figure 4:**
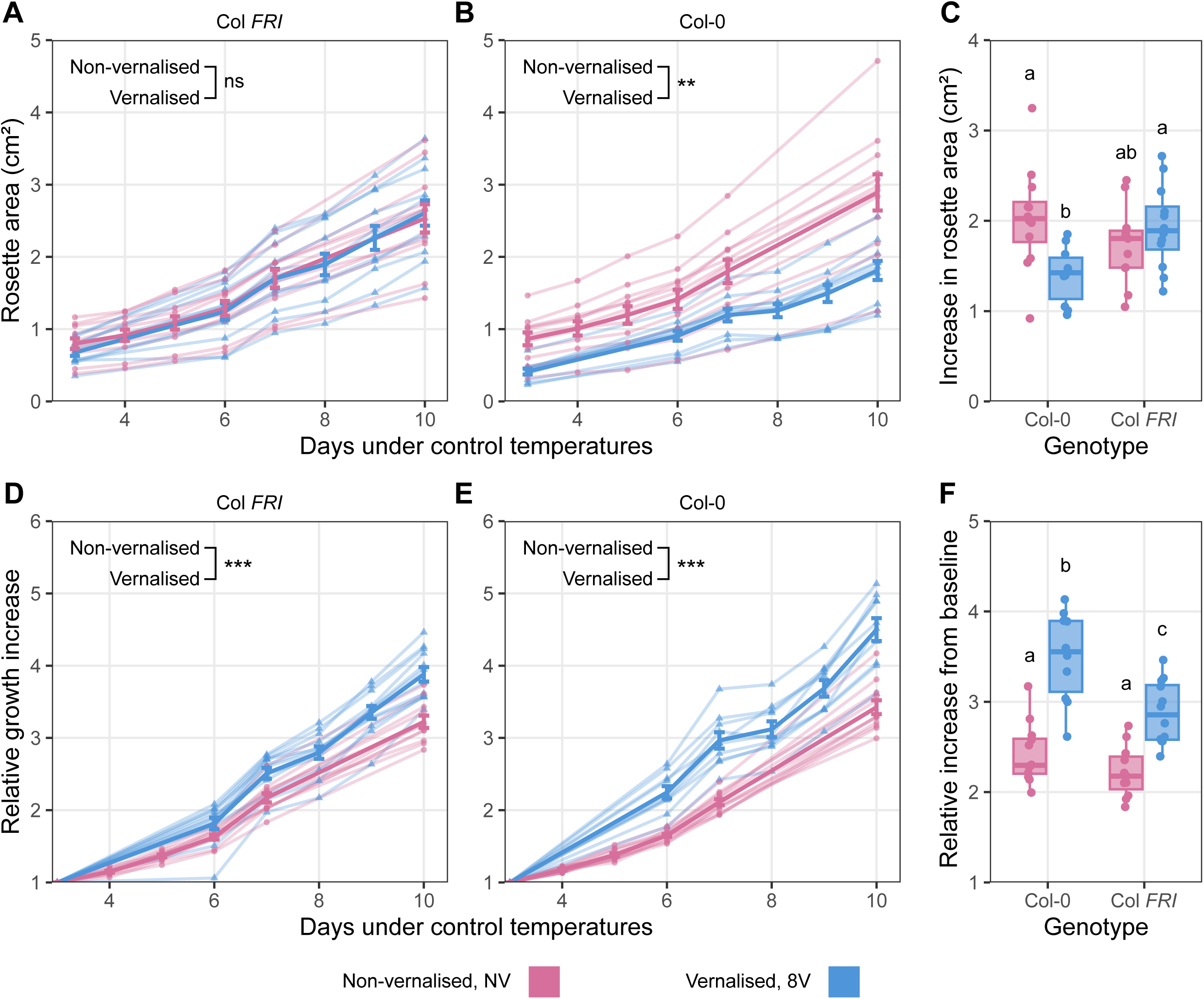
Vernalisation-requiring Col *FRI* and rapid cycling Col-0 have differing growth responses after long-term cold. Arabidopsis seedlings of Col *FRI* and Col-0 were initially grown under control temperature conditions (12L:12D 19 °C) for 11 days before transfer to vernalising cold temperatures (8L:16D, 5 °C) for 8 weeks (8V; NV = not vernalised). Following this, plants were returned to 12L:12D at 19 °C and visible leaf area was monitored. Data shown as absolute rosette area across time of individuals plants for **(A)** Col *FRI* and **(B)** Col-0. Bold line represents mean of all replicates with error bars representing (± s.e.m.). **(C)** Difference between the absolute rosette area of the first and last timepoints. Relative change in rosette size normalised to initial rosette area (on day 3) for **(D)** Col *FRI* and **(E)** Col-0. **(F)** Fold increase of normalized visible rosette size between the first and last timepoints. **(A, B, D, E)** Comparison of growth trajectories was analysed using a linear mixed effects model accounting for repeated measures of individual plants followed by with post hoc Tukey’s HSD test. ns, not significant; * p < 0.05, ** p < 0.01, *** p < 0.001. **(C, F)** Different letters indicate statistically significant differences (p < 0.05) from two-way ANOVA followed by with post hoc Tukey’s HSD test (n = 10-12).

To compare the relative growth rates independently of initial (starting) plant size, the leaf area was normalised to the first measurement at the beginning of the experiment. Using this approach, we could control for differences in growth between the genotypes that occurred previously, during the cold temperature treatment. Overall, vernalised plants grew relatively faster than non-vernalised plants for both genotypes (Fig. 4D-F). This was the case both when comparing treatments using a LMM across the entire time course (Fig. 4D, E; p < 0.001) and when comparing the relative change in rosette size between the start and end the measurements (Fig. 4F; p < 0.001). Under control temperature conditions, there was no difference between the growth rate of Col-0 and Col *FRI* (Fig. 4F; p > 0.5), supporting the analysis of absolute leaf area (Fig. 4C). Comparisons between the growth rates of vernalised plants suggested that effect of vernalisation on growth rate varied between genotypes, such that Col-0 grew faster than Col *FRI* (Fig. 4F).

Together, this suggests that Col-0 plants had smaller rosettes than Col *FRI* after vernalisation, but they may compensate for this though accelerated growth upon return to control temperature conditions. In comparison, Col *FRI* plants achieved a larger rosette the during the vernalisation treatment but have slower growth upon transfer to control temperature conditions.

### Vernalisation-induced alterations in the clock are not determined by FRI or FLC

FRI and FLC are the major determinants of vernalisation requirement in Arabidopsis, and variation in *FLC* silencing is a strong contributor to flowering time post-vernalisation (Napp-Zinn, 1957; Shindo et al., 2005). It has been suggested previously that the response of the circadian clock to vernalisation occurs independently from *FLC* expression (Salathia et al. 2006). To assess the roles of FRI and FLC in this process, we examined the response of the circadian clock to 8 weeks of cold in Col-0 – which lacks a functional *FRI* allele – and *FLClean*, where the entire *FLC* gene was removed by gene editing (Nielsen et al., 2024). To allow comparison of multiple genotypes, we screened two timepoints under free running conditions that corresponded to 1 h after subjective dawn (25 h in constant light) and 1 h before subjective dusk (35 h in constant light). These timepoints were chosen to capture the vernalisation-induced changes in circadian oscillator transcripts that occurred within 48 h time courses (Fig. 1).

Non-vernalised Col *FRI* plants had higher *PRR7* transcript levels at 35 h compared with 25 h (Fig. 5A). In vernalised Col *FRI* plants, *PRR7* transcripts levels did not differ between these timepoints, indicating altered rhythms of *PRR7* transcripts following exposure to prolonged cold (Fig. 5A). This is consistent with earlier experiments (Fig. 1D, H). Similarly, in Col-0, *PRR7* transcripts levels did not differ between the timepoints after vernalisation (Fig. 5B). Loss of *FLC* in *FLClean* also did not impact the effect of vernalisation on *PRR7* dynamics (Fig. 5C). *CCA1* and *TOC1* transcripts responded similarly across the genotypes, demonstrating specificity of the response to certain transcripts within the circadian oscillator (Supplementary Fig. 2A, B). These results suggest that the effects of vernalisation on specific clock transcripts are uncoupled from the roles of *FRI* and *FLC* in the vernalisation.

**Figure 5:**
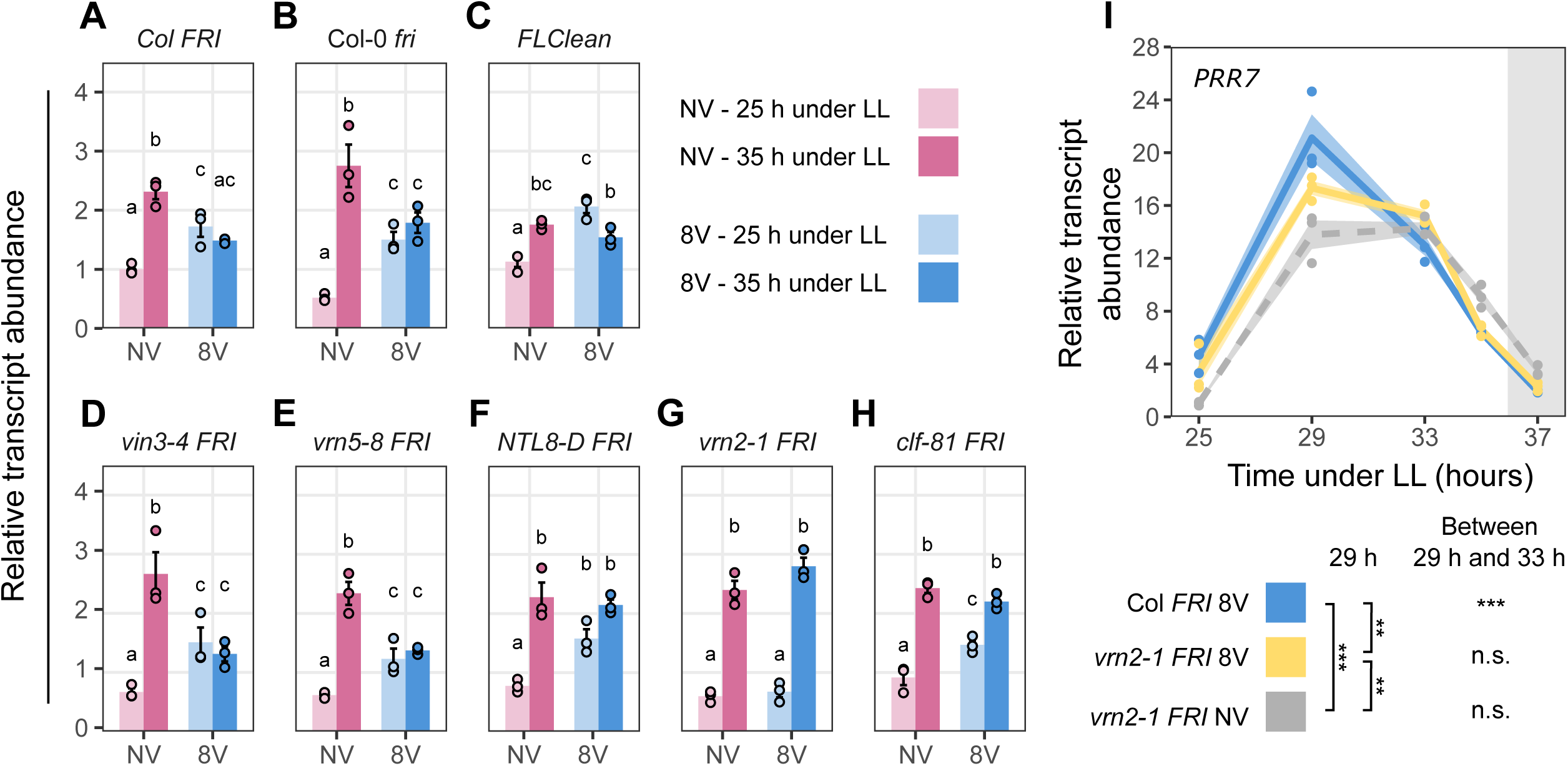
The Polycomb Repressive Complex 2 (PRC2) is necessary for altered dynamics of the circadian oscillator transcript *PRR7* after prolonged cold. Arabidopsis seedlings were grown initially under control temperature conditions (12L:12D 19 °C) for 11 days before transfer to vernalising cold temperatures (8L:16D, 5 °C) for 8 weeks (8V). Non-vernalised (NV) plants were grown for an equivalent amount of developmental time under control conditions. Following this, plants were returned to 12L:12D 19 °C for a further 7 days. On day 8, plants were transferred to constant light conditions (LL). **(A-H)** Relative abundance of key circadian oscillator transcript *PRR7* in following a period of prolonged cold, compared with non-vernalised control plants collected following 25 or 35 hours under constant light. Different letters indicate represent statistically significant differences between transcript levels within each graph (p < 0.05; ANOVA with post-hoc Tukey’s HSD tests for pairwise comparisons). **(I)** Time-series of relative *PRR7* transcript abundance in vernalised Col *FRI* plants (8V, blue), compared with *vrn2-1* plants that were vernalised (8V, yellow) or non-vernalised (NV, grey). Statistical comparisons are between genotypes at following 29 h under constant light and between 29 and 33 hours under constant light, both determined using ANOVA with post-hoc Tukey’s HSD tests for pairwise comparisons. **(A-I)** Transcript levels calculated relative to *PP2AA3*, using RT-qPCR. Data are mean ± s.e.m, n = 3.

### Changes to circadian clock transcripts after vernalisation require the activity of core PRC2 components

Vernalisation depends on the epigenetic silencing of the *FLC* gene. This relies on PRC2 components, including the core proteins VRN2 and CLF, and PRC2 accessory proteins VIN3 and VRN5. Our data suggest that despite entrainment to control temperature conditions, prior exposure to long-term cold is stably registered within the clock. Thus, we set out to determine whether epigenetic regulation contributes to this. We used a variety of PRC2 mutants known to affect *FLC* silencing, and compared the effect of vernalisation on the accumulation of circadian oscillator transcripts.

VIN3 is an early cold-induced component of the vernalisation pathway that is essential for *FLC* silencing (Sung & Amasino, 2004). Therefore, we investigated whether VIN3 contributes to the regulation circadian clock dynamics following prolonged cold. The response of *PRR7* to winter cold was unaltered in *vin3-*4 (Fig. 5D), suggesting that *VIN3* is not required for these responses. VRN5 is a homolog of VIN3 and also required for *FLC* silencing (Greb et al., 2007). We observed that the response of *PRR7* to vernalisation was also unaltered in *vrn5*-8 (Fig. 5E). Temperature-dependent growth, conveyed through NTM1-LIKE 8 (NTL8) protein concentration, also participates in long-term cold sensing through the regulation of *VIN3* (Zhao et al., 2020). However, it remains unclear whether this indirect cold sensing mechanism may regulate a wider set of cold-responsive genes. We observed no changes in *PRR7* transcript levels between vernalised and non-vernalised plants in the gain-of-function *ntl8-D* mutant compared with Col *FRI* (Fig. 5F). In this mutant, *NTL8* is constitutively active in, leading to misregulation of *VIN3* even in the absence of cold (Zhao et al., 2020). Therefore, this further supports our finding that *VIN3* activity is not required for altered circadian clock dynamics following prolonged cold and suggests that this independent cold sensing pathway does not have a broader effect on the circadian clock. There were also no effects of these mutants on post-vernalisation dynamics of *CCA1* and *TOC1* transcripts (Supplementary Fig. 2A, B).

VERNALISATION2 (VRN2) and CURLY LEAF (CLF) are core subunits of the PRC2 and participate in epigenetic silencing of *FLC* and other targets during cold (Wood et al., 2006; Greb et al., 2007). In both *vrn2-1* and *clf-81* mutants, the effect of vernalisation on *PRR7* dynamics is lost (Fig. 5G, H). This indicates that the PRC2, and VRN2 in particular, is required for the response of the circadian clock transcript *PRR7* to winter cold.

To examine further the effect of the *vrn2-1* mutant upon *PRR7* transcript dynamics, we conducted an additional experiment using identical conditions, but sampling at several daytime timepoints when the post-cold phase shift of *PRR7* is greatest under free running conditions (Fig. 5I). In vernalised Col *FRI*, *PRR7* transcript levels were greatest after 29 h of constant light, and lower after 33 h (Fig. 5I). In contrast, in the vernalised *vrn2-1* mutant, *PRR7* transcript abundance was unaltered between 29 h and 33 h of constant light (Fig. 5I). At 29 h, *PRR7* transcript abundance was greater in vernalised Col *FRI* compared with vernalised *vrn2*-1 (Fig. 5I). In combination with the data in Fig. 5G, this indicates that core PRC2 components, particularly VRN2, are necessary for vernalisation-induced alterations in *PRR7* dynamics under free running conditions.

## Discussion

Here, we studied the effect of vernalising cold treatments upon the Arabidopsis circadian clock. We focused on the winter-annual accession Col *FRI*, as we reasoned that its clock might respond differently to these treatments compared with the more commonly used Col-0 rapid cycling accession. The are existing demonstrated links between the circadian clock, the vernalisation pathway, and the floral repressor *FLC* (Edwards et al., 2006; Salathia et al., 2006; Spensley et al., 2009; Kyung et al., 2022). This motivated us to study the effect of vernalisation on the temporal dynamics of specific oscillator components and circadian clock outputs.

We identified that after vernalisation, *PRR7* transcripts in Col *FRI* phased earlier under free running conditions compared with plants grown in control temperatures, despite the same entrainment regime (Fig. 1D). This did not occur for other key clock components such as *CCA1* and *TOC1* (Fig. 1C, E), suggesting differential sensitivity of distinct clock components to environmental factors (Webb et al., 2019; Paajanen et al., 2026). *PRR7* is positioned within the morning loop of the circadian oscillator (Nakamichi et al., 2010; Farré & Liu, 2013). It acts downstream of the phytochrome photoreceptors and participates in temperature compensation mechanisms alongside other *PRR*-family genes (Kaczorowski & Quail, 2003; Salome et al., 2010; Shor et al., 2017; Martín et al., 2018). Thus, it can integrate both light and temperature signals into the clock to regulate oscillator dynamics. Moreover, *PRR7* transcription is influenced by internal metabolic signals communicated *via* the sugar-responsive transcription factor bZIP63 and the energy-sensing kinase SnRK1 (Haydon et al., 2013; Frank et al., 2018). Collectively, this supports the idea that *PRR7* may act as a signalling hub through which both external and internal cues are integrated across the plant (Webb et al., 2019). Our data imply that in addition to this, *PRR7* registers past environmental history through alteration of its phase in response to prolonged cold temperatures (Fig. 1, 5).

In nature, vernalisation ensures that flowering is delayed until plants have been exposed to a sufficient period of cold, preventing premature flowering during inductive photoperiods of autumn. Previous research using high temporal resolution leaf movement assays identified a modest vernalisation-dependent shortening of the Arabidopsis circadian period (up to approximately 1 h shorter) that occurs independently from *FRI* and *FLC* (Salathia et al., 2006). Given that shorter period circadian clock mutants often have early flowering phenotypes (summarised in Wang et al. (2025)), these faster paced clocks have been proposed to support photoperiodic flowering in spring (Salathia et al., 2006). The sampling resolution of our experiments was insufficient to determine whether the same period changes occurred for oscillator transcripts. However, we did observe that vernalisation eliminated the delay in flowering that occurs between long-day and equinox photoperiods, in both Col *FRI* and in Col-0 (Fig. 3B-E). This suggests that prolonged exposure to cold temperature modulates the sensitivity of photoperiodic responses, despite Col-0 not having a vernalisation requirement for flowering. We also identified upregulation of *CO* transcripts at the end of a 12-hour day in vernalised plants (Fig. 2A), which could arise from altered circadian regulation. Since increased amplitude of *CO* expression alters overall *FT* levels and accelerates flowering (Ito et al., 2012), this could contribute to the flowering phenotypes we observed (Fig. 3B).

It was interesting that growth rates were affected by vernalisation (Fig. 4), because growth is also regulated by photoperiod length – albeit through distinct mechanisms (Liu et al., 2021; Wang et al., 2024). The metabolic daylength measurement system that underpins seasonal growth regulation is based on combined circadian clock- and photoperiod-controlled regulation of starch degradation (Liu et al., 2021; Li & Gendron, 2025). *MYO-INOSITOL-1-PHOSPHATE SYNTHASE 1* (*MIPS1*) is required to support leaf growth under long days, with a critical photoperiod for the light-sensitive phase for *MIPS1* expression to occurring after ZT12 in long-days (Wang et al., 2024). We speculate that the same winter cold-induced change in circadian dynamics could also extend to other circadian outputs, including *MIPS1* expression. If so, this could change the critical photoperiod for growth, thereby explaining why vernalised plants grow relatively more than non-vernalised plants.

By disrupting components of the vernalisation pathway, we identified that the PRC2 subunit VRN2 is required for the cold-induced phase shift of *PRR7* transcripts (Fig. 5A, E). *VRN2* mRNA is expressed constitutively and its protein is stabilised under prolonged cold (Gibbs et al., 2018). This suggests a potential link between proteins that form part of the vernalisation pathway and circadian clock function (Salathia et al., 2006). In Arabidopsis, *VRN2* is also associated with seed dormancy (Auge et al., 2017) and development (Roszak & Köhler, 2011), root architecture (de Lucas et al., 2016), hypoxia stress responses (Labandera et al., 2021), leaf growth (Osborne et al., 2025), and flowering under short days (Labandera et al., 2021). Evidence for a direct interaction between the circadian clock and VRN2 remains limited (Xi et al., 2020; Osborne et al., 2025). One explanation is that the interaction may be context-dependent and detectable only following cold exposure. Alternatively, the relationship between VRN2 and the clock may be mediated indirectly by unknown intermediate factors. Through exploration of public datasets, we generated a list of 771 potential candidate genes that could influence *PRR7* dynamics following vernalisation (Supplementary Dataset 2). These genes are stably down-regulated during vernalisation, as is expected if they are repressed through the action of VRN2-PRC2 during cold (Xi et al., 2020), but are not targets of *VIN3* or *VRN5* (Franco-Echevarría et al., 2023). Interestingly, this change in *PRR7* dynamics appears to be FLC-independent. This is consistent with post-cold *PRR7* dynamics being unaltered by the absence of *FLC* expression in Col-0 and *FLClean*. Moreover, loss of VIN3 or VRN5 – which have critical roles in *FLC* silencing – had no effect on the vernalisation-induced response of *PRR7*.

Our data suggest that following vernalisation, there are adjustments to the circadian clock. This might help to align whole-plant physiology with the springtime environment. It could potentially extend to other circadian regulated processes, ensuring they are regulated appropriately for the season. For example, the phase of the ‘autumn clock’ may favour processes associated with resource allocation and cold tolerance. On the other hand, the ‘spring clock’ may favour accelerated growth following winter dormancy and reinforce the reproductive transition. Although we focused on Arabidopsis, vernalisation is part of the seasonal life cycle in many other plant species (Li et al., 2013; Wu et al., 2015; Bouché et al., 2017; O’Neill et al., 2019). Therefore, our work could provide insights into the regulation of seasonal responses in plants including those that hold agronomic importance, such as other Brassicas and cereal crops.

## Materials and methods

### Plant material and growth conditions

Background accessions of *Arabidopsis thaliana* (Arabidopsis) that were used were Columbia-0 (Col-0) and Col *FRI^sf2^* (gifted from Caroline Dean at the John Innes Centre (Lee et al., 1994)). All mutant germplasm was in the Col *FRI* background and donated by Caroline Dean (John Innes Centre). These were *vin3-4,* which contains a T-DNA in the second intron of *VIN3* (Bond et al., 2009); *flc-2,* a fast-neutron allele containing a 30 kb deletion spanning the *FLC* locus (Michaels & Amasino, 1999); *FLClean,* a CRISPR-Cas9 deletion of the entire *FLC* genomic sequence (Nielsen et al., 2024); *vrn2-1* (Yang et al., 2017); *vrn5-8,* a T-DNA insertion line (SALK_136506) (Greb et al., 2007) crossed into the *FRI* background; *clf-81*, an EMS line with a point mutation (Kim et al., 1998) crossed into the *FRI* background, and *ntl8-D3*, a truncated NTL8 protein caused by a deletion in the C-terminal transmembrane domain resulting in enhanced nuclear localisation (Zhao et al., 2021).

For cultivation, Arabidopsis seeds were surface sterilised using 70% (v/v) ethanol for 10 minutes with agitation, washed 3 times with sterile water, and suspended in 0.1% (w/v) agar. Sterilised seeds were pipetted onto half strength Murashige and Skoog (MS) media with 0.8% (w/v) agar (pH 6.8), without added sucrose, and stratified in darkness at 4°C for 3 days before being moved into growth chambers. Cultivation occurred in PHCBi MLR-352 growth chambers.

### Circadian time-course sampling for RT-qPCR

Following stratification, seeds were grown under controlled temperature conditions (19 °C, 12L:12D, ∼ 50 μmol m^−2^ s^−1^) for 11 days. On day 12 after transfer to growth chambers, plants were moved to vernalisation conditions (5°C, 8L:16D, ∼ 20 μmol m^−2^ s^−1^) for 8 weeks. Following this, to investigate persistent changes within the circadian oscillator, plants were initially returned to control temperature growth conditions (19 °C, 12L:12D, ∼ 50 μmol m^−2^ s^−1^) for 7 days. On day 8 of this condition, to monitor the circadian oscillator under free running conditions, plants were transferred to constant light and temperature. Sampling commenced after 24 h of constant light to avoid sampling during transitory effects of the final dawn. To control for potential effects of developmental age on circadian clock gene expression (Kim et al., 2016), non-vernalised controls were grown under control temperature conditions for an amount of developmental time that was equivalent that elapsed under the vernalising conditions, using an approach described previously (Zhao et al., 2020). Where possible, the growth of non-vernalised seedlings was staggered such that sampling occurred on the same day as vernalised seedlings.

For 48-hour time course sampling, after 24 hours of constant light, aerial plant tissue was collected every 4 hours for 48 h (from 25 h to 73 h of constant light). For experiments investigating vernalisation pathway mutants, after 24 hours of constant light, aerial plant tissue was collected after 25 h and 35 h of constant light, as these timepoints allowed efficient discrimination of effects of the vernalisation mutants upon circadian clock gene expression across multiple mutant backgrounds. To gain a more detailed understanding of circadian clock transcript dynamics in the *vrn2-1* mutant following 8 weeks of vernalisation, plants were grown as described previously and samples were collected under constant light conditions after 25 h, 29 h, 33 h, 35 h and 37 h of constant light. For each time point, aerial tissue from five individual plants was pooled to generate one biological replicate. A total of three biological replicates were obtained per time point. Harvesting was destructive, precluding repeated measurements on the same plant. Tissue was immediately frozen in liquid N_2_ and stored at −70 °C until processing.

### RNA isolation and RT-qPCR

Total RNA from Arabidopsis seedlings was prepared using RNeasy Plant MINI kits (Qiagen) with on-column DNase treatment using the RNAse-Free DNase Set (Qiagen), as per manufacturer’s instructions. RNA was eluted using nuclease-free water and sample quality was assessed (Nanodrop, Thermo-Scientific). RNA was stored at −70°C until required.

For RT-qPCR analysis, 1 µg of total RNA was used to synthesise cDNA using a High-Capacity cDNA Reverse Transcription kit following manufacturer’s instructions, using random primers (Applied Biosystems, ThermoFisher). Relative transcript abundance was measured using a LightCycler 480 II (Roche), using four technical replicates per sample and cDNA that was diluted 1:100 using nuclease-free water. *PROTEIN PHOSPHATASE 2A SUBUNIT A3* (*PP2AA3*) was used as a refence transcript as its expression is arhythmic (Supplementary Fig. 3) and appears unaltered by cold treatments (as determined by EFP browser, Waese et al., 2017). Total reaction volume was 10 µl, containing: 5 µl 2x RT-qPCR BIO SyGreen LO-ROX (PCR Biosystems); 2.5 µl template cDNA; 0.4 µl 10 mM forward primer and 0.4 µl 10 mM reverse primer. Primers used are detailed in Supplementary Table 1. Absolute quantification of Ct values was determined using the Second Derivative Maximum method (Guescini et al., 2008). Relative transcript abundance was calculated using the 2^ΔΔCt method and normalised to Col FRI NV at 25 h in constant light.

### Circadian time-course data analysis

For RT-qPCR time-series spanning 48 h, data were analysed with several methods that are used to examine properties of oscillatory data, including JTK_CYCLE (Hughes et al., 2010) and meta2d within the R package MetaCycle, using default parameters (Wu et al., 2016). Fast Fourier Transform-Non Linear Least Squares Method (FFT-NLLS) used BioDare2, with no detrending (biodare2.ed.ac.uk, Zielinski et al., 2014), and CircaCompare was used when the period length was approximately 24 h (Parsons et al., 2020). Phase was reported in circadian units, i.e. using time-series aligned to a 24 h period (sometimes known as “modulo tau”). Changes in phase, period and amplitude following vernalisation were calculated by taking the mean of these parameters for 8-week vernalisation treated samples and subtracting this from these parameters from the non-vernalised dataset.

To examine the transcript abundance of circadian clock outputs, RT-qPCR was performed on samples collected between 25 h and 49 hours under constant light normalised to the refence transcripts *PP2AA3* and *UBQ10*. Data were analysed at each time point between vernalisation treatments using Welch’s two-sample t-tests. To account for multiple testing, p-values were adjusted using the Benjamini–Hochberg false discovery rate (FDR) procedure.

### Growth assays

To investigate plant growth dynamics under control temperatures, following vernalisation, Col-0 and Col *FRI* seeds were sown on 0.5 MS agar and stratified. They were initially grown under control temperature conditions (19 °C, 12L:12D, ∼ 50 μmol m^−2^ s^−1^) for a minimum of 11 days, then transferred to vernalising conditions for 8 weeks (5°C 8L:16D, ∼ 20 μmol m^−2^ s^−1^). Following this, 12 plants were transplanted to compost (Levington F2 starter mix) in 24-compartment horticultural trays and returned warm control conditions for 10 days.

Germination and growth of non-vernalised seedlings was deliberately delayed so that these plants were transferred to compost at the equivalent developmental stage as vernalised seedlings, to allow side by side comparison (Zhao et al., 2020).

During the period following prolonged cold, plants were imaged daily at approximately the same time of day using a camera positioned directly above the compost trays. A calibration scale was always included in images. Visible leaf area was measured using FIJI software (Schneider et al., 2012). Using this, images were manually thresholded so the entire rosette was isolated from the compost background. Thresholded images were converted to binary format, allowing measurement of the area of each thresholded region.

### Flowering time assays

Seedlings were grown initially on 0.5 MS agar under short day conditions (8L:16D) and 19°C, to prevent photoperiodic transition to flowering. After this growth period (minimum of 11 days), plants were transferred to vernalising conditions (5°C 8L:16D, ∼ 20 μmol m^−2^ s^−1^) for either 0, 4, or 8 weeks. Plants receiving the NV or 4-week vernalisation treatments (*vs.* full 8 week vernalisation) were given additional growing days under warm short days to account for developmental differences incurred during the cold period (Zhao et al., 2020). In this experiment, the growth of non-vernalised seedlings was not staggered relative to the vernalised seedlings, and plants were transferred to compost at equivalent developmental stages but on different days. Following vernalisation, plants were transplanted onto compost (Levington F2 starter mix) and transferred to control temperatures (19°C) under 8L:16D, 12L:12D or 16L:8D photoperiods. These were chosen as standard short day, equinox or long day protocols with Arabidopsis. Plants were monitored regularly until bolting, and flowering time was scored as below.

Flowering was classified as the day at which the terminal inflorescence bud exceeded 10 mm. Total leaf number (TLN) was also measured, to estimate the developmental stage at flowering. The TLN included all primary leaves in the rosette, including cotyledons, but excluded cauline and auxiliary leaves. The total number of days under control conditions following vernalisation was also recorded, to calculate days to bolting (DTB). For comparisons of TLN and DTB with genotypes, two-way ANOVA was used with vernalisation treatment length and photoperiod as interacting factors P-values were adjusted using the Tukey’s HSD test for post-hoc contrasts.

### Analysis of growth dynamics

To compare differences in visible leaf area (VLA) between genotypes and growth conditions at specific time points, two-way ANOVA was performed with genotype and growth condition as fixed factors. Post-hoc pairwise contrasts were conducted using Tukey’s HSD test to correct for multiple testing. Relative leaf area was calculated as follows (day 3 was the first day of imaging):

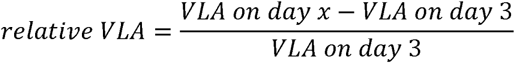

To compare growth over time, repeated measurements of both absolute and relative VLA were analysed using linear mixed models (LMM) using the lmerTest package (Kuznetsova et al., 2017). Because each plant was measured at multiple time points, plant identity was modelled as a random factor to account for the non-independence of repeated observations. Area was modelled as a function of time, genotype, treatment, and their interactions. To account for the uncertainty in estimating random effects, P-values for fixed effects were obtained using Satterthwaites approximation using the lmerTest package (Kuznetsova et al., 2017). Pairwise contrasts were conducted using the emtrends function in the emmeans package (Lenth et al., 2025), with Tukey’s adjusted p-values for multiple testing:

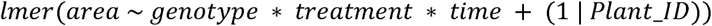

### Statistical analysis

All statistical analyses and graphing was performed using R v4.4.1.

## Acknowledgements

This work was funded by UKRI-BBSRC (Institute Strategic Programmes GEN BB/P013511/1 and BRiC BB/X01102X/1), the John Innes Centre, and the John Innes Foundation. MM is funded by Wellcome Trust Early-Career Award 325529/Z/25/Z. We thank Maximillian Jones for comments on the manuscript, Caroline Dean for seed donation, and Anna Schulten, Steve Penfield, and Martin Howard for scientific advice.

## Data availability

Source data for this study are in Supplementary Dataset 3.

## Supplementary Datasets

**Supplementary Dataset 1.** Quantification of oscillations of circadian clock transcripts before and after vernalisation. Estimates of rhythmic parameters of circadian oscillator components in non-vernalised (NV) and vernalised (8V) Col *FRI* plants entrained under 19 °C 12L:12D released into free-running conditions.

**Supplementary Dataset 2:** Potential candidate genes that may mediate the vernalisation dependent phase shift of *PRR7* transcripts in Col *FRI*.

**Supplementary Dataset 3:** Source data for this study.

