## Supplementary Figures and Table for "Vernalisation-induced changes to the Arabidopsis circadian clock require Polycomb Repressive Complex 2 and are FLC-independent"

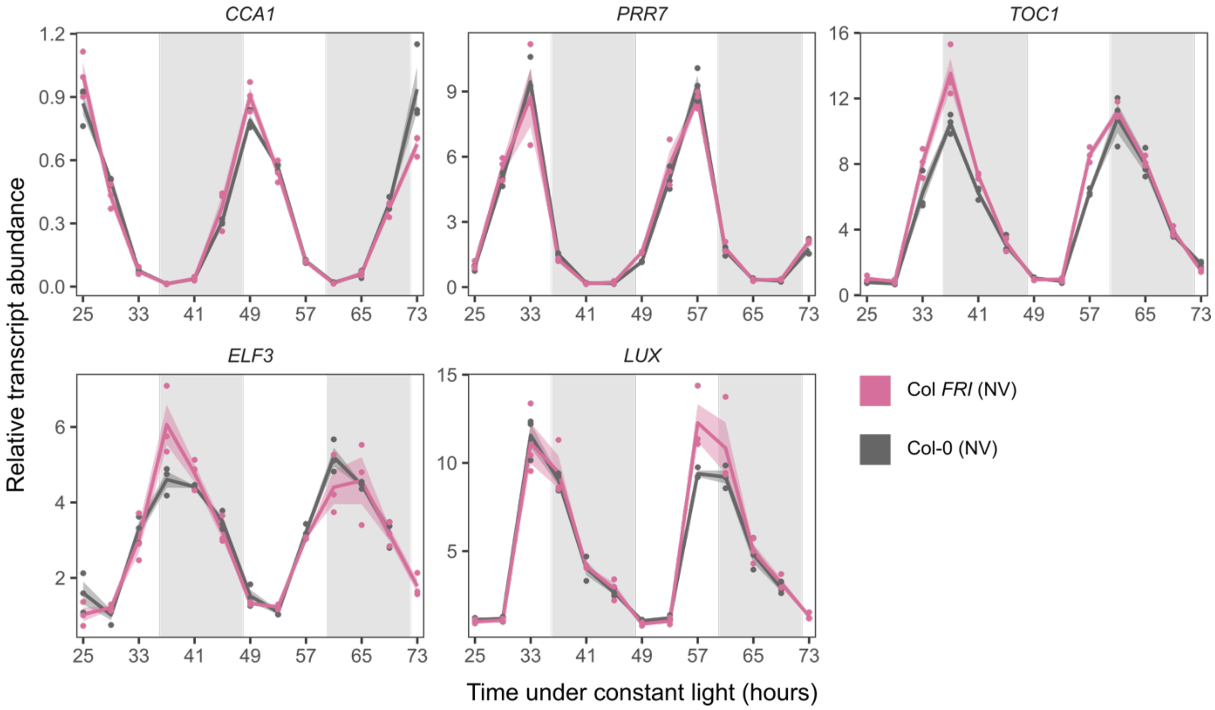


**Supplementary Figure 1.** Transcript dynamics of circadian oscillator components in non-vernalised Col FRI and Col-0 plants. Relative transcript abundance of circadian oscillator transcripts *CCA1*, *PRR7*, *TOC1*, *ELF3* and *LUX* in non-vernalised Col *FRI* and Col-0 plants. Arabidopsis seedlings were grown under control temperature conditions (12L:12D 19 ˚C) for 26 before being transferred to constant light conditions. Sampling of aerial tissue for RNA started after 25 h of constant light with tissue collected every 4 hours for 48 hours. For each timepoint, aerial tissue from up to five individual plants was pooled to generate each biological replicate. Solid lines indicate mean, with shaded ribbon ± s.e.m (n = 3). Grey shading indicates subjective night. NV = not vernalised.

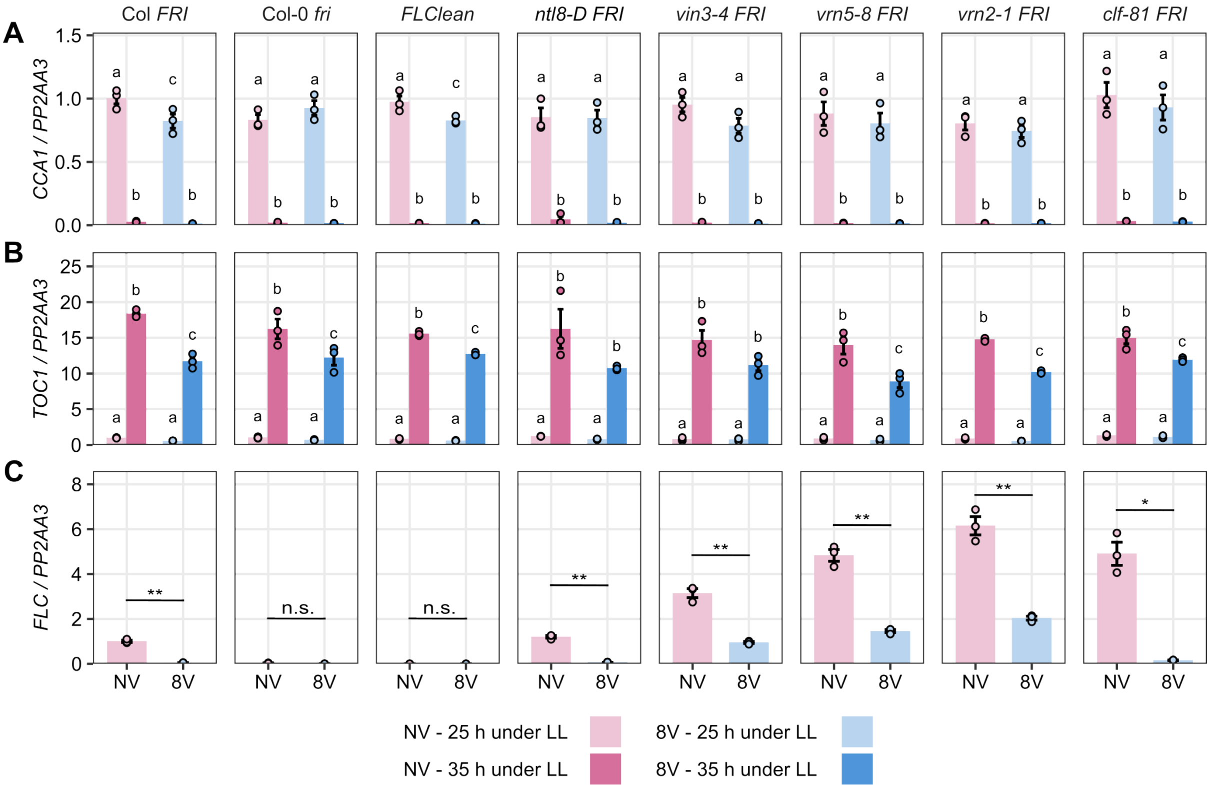


**Supplementary Figure 2:** Effect of vernalisation pathway mutants upon *CCA1* and *TOC1* transcript levels. **(A, B)** Relative abundance of key circadian oscillator transcripts *CCA1* and *TOC1* in following a period of prolonged cold (8V), compared with non-vernalised (NV) control plants collected following 25 or 35 hours under constant light (LL). Different letters indicate statistically significant differences between transcript levels within each plot ($p < 0.05$; ANOVA with post-hoc Tukey's HSD tests for pairwise comparisons). **(C)** Relative abundance of *FLC* transcripts in NV and 8V plants measured at after 25 h under constant light. Asterisks indicate statiscal comparison between vernalised and non-vernalised plants within each genotype (Student's t-test; * $p < 0.05$, ** $p < 0.01$, n.s. $p >= 0.05$). Transcript levels calculated relative to *PP2AA3*, using RT-qPCR. Data are mean ± s.e.m, n = 3.

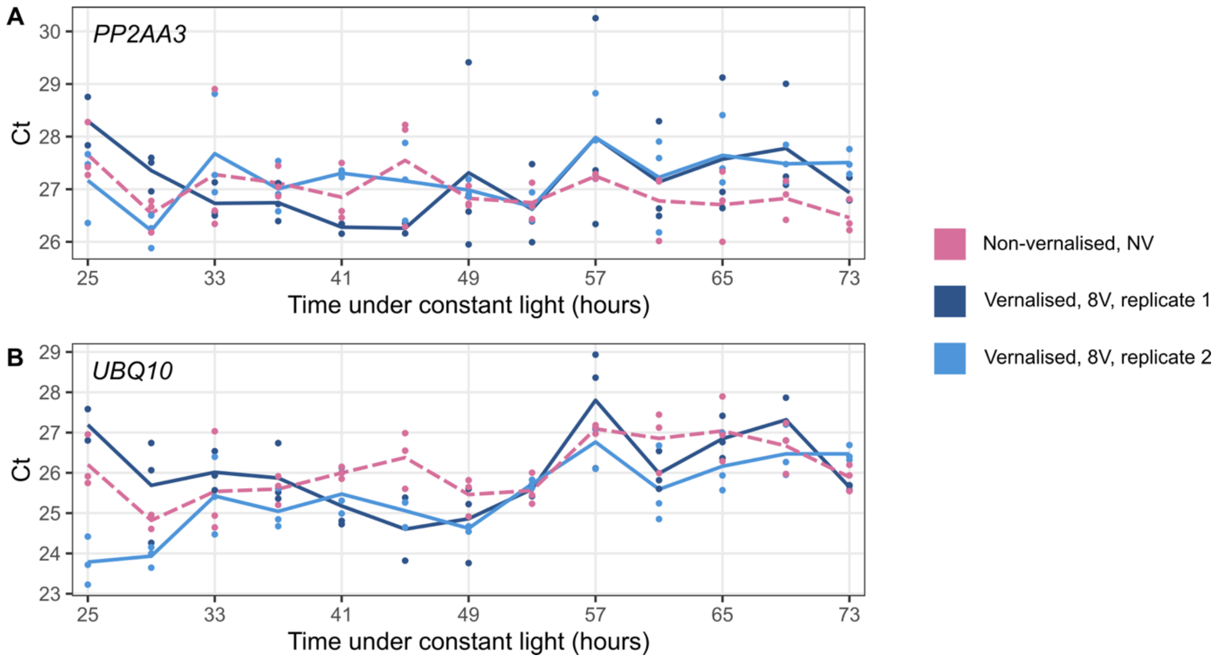


**Supplementary Figure 3.** Inspection of reference transcripts. Reference transcripts **(A)** *PP2AA3* and **(B)** *UBQ10* were not rhythmic during time courses in our experiments, determined by JTK_CYCLE ($p > 0.5$). Line indicates mean Cp value ($n = 2\text{-}3$).

| Primer | Purpose | Primer sequence (5’ – 3’) |
|---|---|---|
| ABA1-F | RT-qPCR | AGAGCAACACGGAATTTTCC |
| ABA1-R | RT-qPCR | TCGATTTCGGAGTTTTCCTG |
| CBF1-F | RT-qPCR | CCGCCGTCTGTTCAATGGAATCAT |
| CBF1-R | RT-qPCR | TCCAAAGCGACACGTCACCATCTC |
| CBF2-F | RT-qPCR | AACTCCGGTAAGTGGGTGTG |
| CBF2-R | RT-qPCR | CGGCGTATAAATAGCCTCCA |
| CCA1-F | RT-qPCR | TCTGTGTCTGACGAGGGTCG |
| CCA1-R | RT-qPCR | ACTTTGCGGCAATACCTCTCTGG |
| CO-F | RT-qPCR | CTACAACGACAATGGTTCCATTAC |
| CO-R | RT-qPCR | CAGGGTCAGGTTGTTGC |
| ELF3-F | RT-qPCR | ACCGAGATGGTGGCAAAACT |
| ELF3-R | RT-qPCR | ACTGCCATGACCCTCTTGTG |
| FER3-F | RT-qPCR | TCCAAGAACGATGATGTCCA |
| FER3-R | RT-qPCR | CCATGTCCTTTGCCAAGTCT |
| FKF1-F | RT-qPCR | GGATTGTACAACCTTCCCGC |
| FKF1-R | RT-qPCR | ACCCCTTCCCCACCAAATAG |
| HY5-F | RT-qPCR | GTCATCAAGCTCTGCTCCACAT |
| HY5-R | RT-qPCR | CGGCACTCGCCGTATCTC |
| LUX-F | RT-qPCR | TAACGTGGAGGAGGAAGATCGA |
| LUX-R | RT-qPCR | TCCATCACCGTTTGATGTCTTT |
| PIF4-F | RT-qPCR | GCCGATGGAGATGTTGAGAT |
| PIF4-R | RT-qPCR | CCAACCTAGTGGTCCAAACG |
| PIF5-F | RT-qPCR | AAAACCCGGTACAGTTGCAG |
| PIF5-R | RT-qPCR | ATCCAAATCCCAACATGTCC |
| PP2AA3-F | RT-qPCR | CGTTACTGCCAGCCATTGTA |
| PP2AA3-R | RT-qPCR | GCAAAGAGCACCAAGCTTCT |
| PRR7-F | RT-qPCR | CCACGAGCGGTATCTCTATGG |
| PRR7-R | RT-qPCR | ACTTGGAAACTCAGGGTTAGAATGTAC |
| PRR9-F | RT-qPCR | AGCATGAAGCTTGCAAGAAC |
| PRR9-R | RT-qPCR | GCAGCACCTCTCAGCATACA |
| RVE2-F | RT-qPCR | TCCTTTCGAATGAAATGCTTG |

| | | |
|---|---|---|
| RVE2-R | RT-qPCR | TCTATGGCAGAGCTTGGAGAC |
| STO-F | RT-qPCR | CCATCCACGTGGCTAATTCT |
| STO-R | RT-qPCR | AGCCTTCTGTTGGTTGTTGG |
| TOC1-F | RT-qPCR | TCTTCGCAGAATCCCTGTGAT |
| TOC1-R | RT-qPCR | GCTGCACCTAGCTTCAAGCA |
| UBQ10-F | RT-qPCR | GGCCTTGTATAATCCCTGATGAATAAG |
| UBQ10-R | RT-qPCR | AAAGAGATAACAGGAACGGAAACATAGT |

**Supplementary Table 1:** Primers used in this study.
